# Limitations of Solar-Induced Chlorophyll Fluorescence (SIF) for Estimating Photosynthesis Under Stress

**DOI:** 10.1101/2022.11.09.515325

**Authors:** Amir M. Mayo, Menachem Moshelion, Oded Liran

## Abstract

High-throughput measurements of photosynthesis of plants grown under various conditions may provide important insights into the plasticity of the photosynthetic performance of plants. Remote sensing of photosynthetic activity is the next generation of fast scanning techniques, enabling high-throughput photosynthesis measurements under controlled conditions. We hypothesized that by measuring SIF simultaneously with whole-plant water relations in a standardized controlled drought experiment, we would be able to quantify photosynthetic activity and to detect water stress at an early stage. A functional-phenotyping platform was used to apply the controlled drought treatment and to monitor the growth and water balance of tomato introgression lines (ILs). A new SIF-derived index, electron transport rate (RS-ETRi), was found to be negatively correlated with whole-plant stomatal conductance (Gsc) under non-stressed conditions. No significant relationships were found between SIF and plant biomass or Gsc. SIF_687_ responded to drought earlier than any of the other measured vegetation indices. SIF based indices could not differentiate between introgressed lines of tomato; whereas differences between Introgression Lines were clearly identified by the water-relations measurements. We concluded that SIF did not provide any advantage over commonly used methods for detecting physiological differences between the Introgression Lines. Overall, although SIF plays a significant role in photosynthesis, the relationship between SIF and photosynthesis is complex and we believe it would be an oversimplification to use SIF to quantify photosynthetic activity on close canopy spatial resolution level.

## Introduction

Ensuring food security for a growing global population is an unquestionable challenge. The past century has seen substantial growth in food production, largely due to the industrial and agricultural revolutions and, more recently, major advances in molecular and genomic tools, which have revolutionized plant breeding (Pérez-De-Castro et al., 2012). Nevertheless, food producers are experiencing greater competition for land, water and energy, all as they face the threat of substantial climate change and the negative effects of food production on the environment (Tilman et al., 2001). One of the most pressing challenges humanity faces is increasing drought events. Drought is a meteorological term that is commonly defined as a period without significant rainfall and considered to be a major bottleneck for global agricultural productivity around the world. Generally, drought stress occurs when the available water in the soil is reduced and atmospheric conditions cause continuous loss of water by transpiration or evaporation (Jaleel et al., 2009). One way to sense if the plant is found in water stress is to check its photosynthetic rate. It is widely accepted that there is a strong relationship between stomatal conductance and photosynthesis (Farquhar & Sharkey, 1982; S. C. Wong et al., 1979), as the stomata control the insertion of CO_2_ into the leaf. A wide array of studies that have reported on the linear correlation between gas exchange, CO_2_ assimilation (A), and stomatal conductance (g or gs) can be found in the literature (Field, 1987; Radin et al., 1988; S.-C. Wong et al., 1985; S. C. Wong et al., 1979). Additional to carbon assimilation measurements, chlorophyll fluorescence is another signal which is interpreted as photosynthetic activity. Information about the efficiency of photochemistry and Non-Photochemical Quenching (NPQ) can be obtained by measuring the yield of chlorophyll fluorescence (Chlf) (Baker, 2008; Genty et al., 1989).

Light energy that is absorbed by the leaf’s photosynthetic apparatus can be funneled into one of three pathways: It can be used to drive photosynthesis (photochemistry), it can be dissipated via the regulated processes of NPQ or it can be re-emitted as fluorescence. These three processes compete for the same pool of absorbed energy, such that any increase in the quantity of one will result in a decrease in the yield of the other two (Maxwell & Johnson, 2000). ChlF theory and its practical application has been investigated for years on a spatial leaf level (Porcar-Castell et al., 2014). A new wave of developments enables the measurement of ChlF from remote-sensing platforms (Porcar-Castell et al., 2014). The remote sensing of ChlF is based on the passive acquisition of Solar-Induced chlorophyll Fluorescence (SIF, sometimes referred to as sun-induced chlorophyll fluorescence; Plascyk, 1975). SIF was shown to correlate well with gross primary production as long as it is measured on a very low spatial scale resolution (Guanter et al., 2014). As spatial resolution increases there is a dissociation of the SIF from primary production (Marrs et al., 2020). Liran et al. (2020) suggested a new spectral index which is based on SIF and correlates with electron transport rate of the light reactions of plants in order to bypass the high spatial resolution dissociation problem. Further, Liran (2022) shows that this index – Remote Sensing of Electron Transport Rate (RS-ETR) interprets the fluorescence emission as Light Use Efficiency of the photosynthetic apparatus of the crop that it measures, thus relating photochemistry to SIF properties.

only few works have tried to link SIF with water status-related traits (Lu et al., 2018; Xu et al., 2018). Several studies have investigated the response of SIF to water stress and the predominant findings suggest that reduced photosynthetic efficiency and increased NPQ lead to a decline in the emission of fluorescence in response to water stress (Ač et al., 2015; Dobrowski et al., 2005; Xu et al., 2018).

Novel high-throughput selection systems have been developed to enable the rapid screening of large plant populations in controlled, pre-field environments (Halperin et al., 2017). SIF and SIF based indices can become an additional tool that reports on valuable information regarding photosynthetic traits of each sample during such a screen. This information, in conjunction with a high-throughput system output, such as transpiration rate, canopy stomatal conductance (Gsc) and growth rate at high spatiotemporal resolution, will reveal new information about the relationship between SIF and the functional response of plants to natural as well as stress conditions. In this study we employ SIF measurement technique on top of a high-throughput parameters acquisition, on the background of various genotypes of tomato (*S. licopersicum*) that experience drought conditions.

## Results

### Construction of a high throughput - SIF phenotyping platform

Two highly spectral sensitive spectroradiometers (**Figure 1A**), were mounted onto a custom-made cart with an adjustable arm (**Figure 1B**). The platform was moving over tomato plants which some of them were treated with two PSII inhibitors (i.e., 3-(3,4-dichlorophenyl)-1,1-dimethylurea (Diuron) and 4-Amino-3-methyl-6-phenyl-1,2,4-triazin-5-one (Brevis)). Significant differences in fluorescence emission were detected between the control group which did not receive photosynthetic inhibitors and both treatment groups within and between dates (**Figure 1C**). Normalized Differential Vegetation Index (NDVI) which was used as a control for correct growth showed a significant opposite trend, as expected (**Figure 1D**).

**Figure 1.**
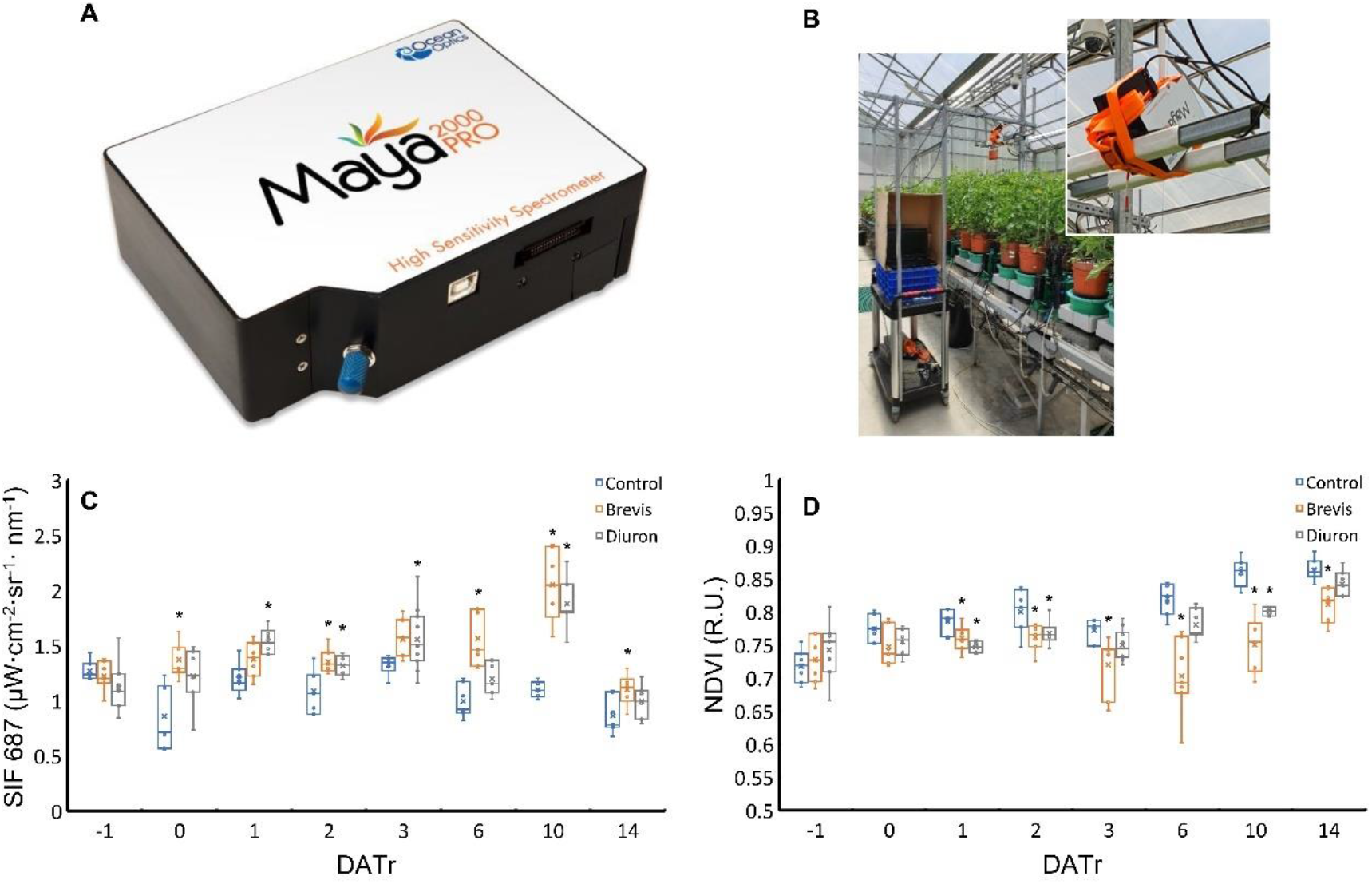
Setup of the SIF-Phenomics platform. (**A**) High-spectral sensitive spectroradiometer (Maya 2000 Pro, Ocean Insight, FL, USA). (**B**) The sensor custom-made setup in the greenhouse. (**C**) Temporal track of SIF_687_ response to photosynthetic inhibitors treatment. (**D**) Temporal track of NDVI response to photosynthetic inhibitors treatment. DATr = days after treatment. Diuron and Brevis were sprayed on the plants at 07:00, DATr 0. Data acquisition was performed around 11:30 - 12:00 each day. 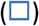 Control group; 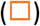 Brevis-treatment group; 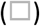 Diuron-treatment group. Asterisk (*) indicates a significant difference between the treatment groups and control group on the same day (Dunnett’s test, *P <* 0.05, 7 ≤*n ≥* 11).

We were interested to compare the correlation between classic reflectance-based Vegetation Indices (VIs) and SIF based indices with chemically extracted photosynthetic pigments concentration along the experimental period. (**Figure 2**)

**Figure 2.**
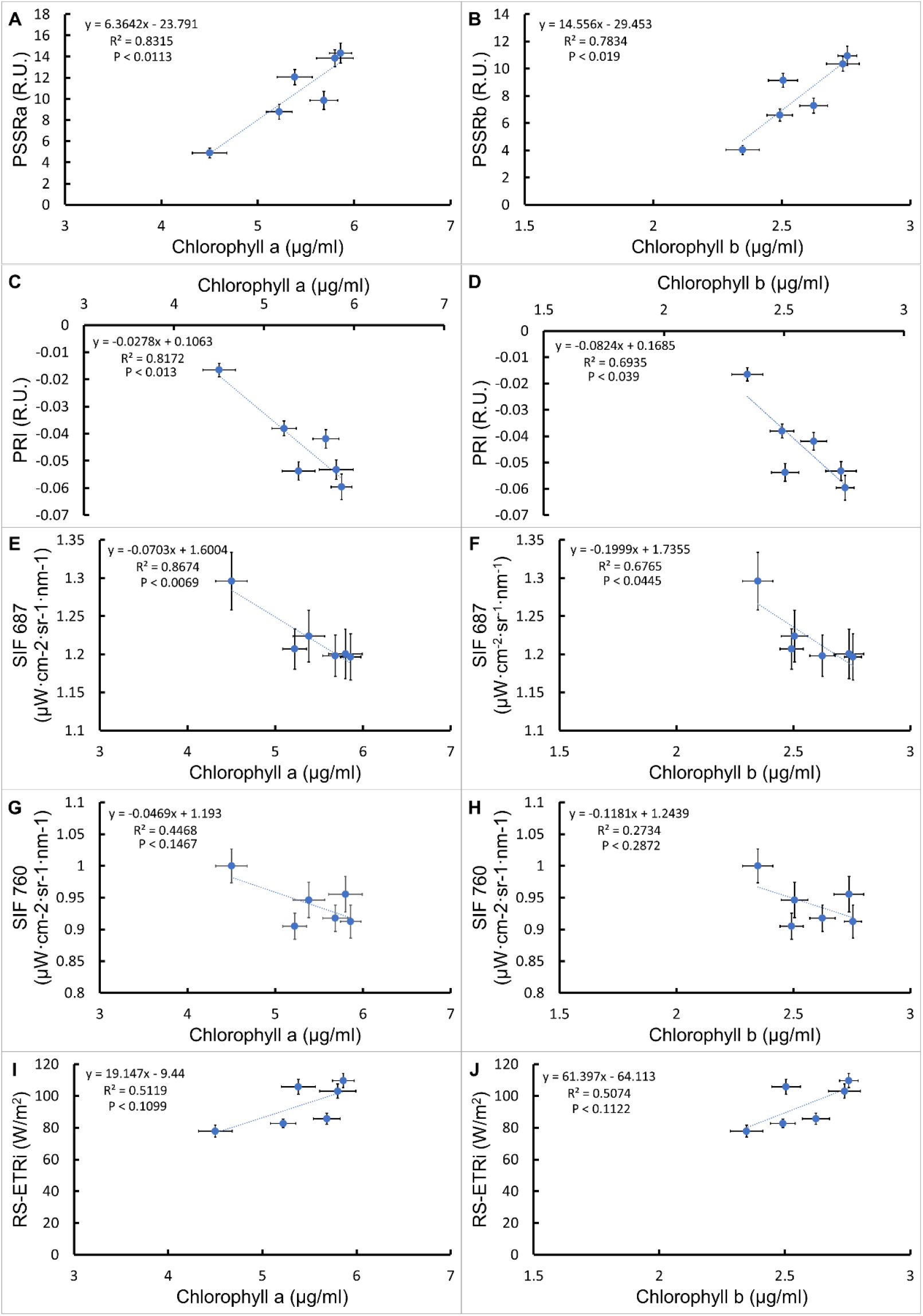
Relationship between leaf pigments concentrations and Vegetation Indices. Correlation analysis between PSSRa, PSSRb, PRI, SIF_687_, SIF_760_ and RS-ETRi to chlorophyll a and chlorophyll b in panels (A)-(J), respectively. Each point is a mean of 20–25 biological replicates ± SE. Values are means of measurements from three different experimental phases: pre-drought, drought and recovery.

Pigment Specific Simple Ratio a (PSSRa) and PSSRb were significantly correlated with the concentrations of Chlorophyll a (Chl a) and Chlorophyll b (Chl b), respectively (**Figure 2A and 2B)**. Significant negative linear correlations were observed between Photosynthetic Response Index (PRI) and the chlorophylls extractions (**Figure 2C, 2D**). SIF_687_, but not SIF_760_ showed significant negative linear correlations to the chlorophyll’s extraction **(Figure 2E-2H)**. Positive non-significant trends were observed between RS-ETRi and pigment concentrations (**Figure 2I, 2J**).

We also checked how well SIF based indices correlate with the high throughput platform outputs – Canopy conductance (Gsc) and net plant weight (**Figure 3**). SIF_760_ and SIF_687_ showed non-significant negative trend with increase in canopy conductance during stress and recovery (**Figure 3A and 3C**, orange dots). They presented a non-significant trend when plants left to grow without stress periods (**Figure 3A, 3C**, blue dots). Electron transport rate (RS-ETRi) showed opposite significant correlations with canopy conductance to those of the fluorescence emissions (**Figure 3E**). Both SIF_687_ and SIF_760_ did not correlate at all with net plant weight (**Figure 3B, 3D**), whereas electron transport rate (RS-ETRi) correlates logarithmically and significantly with increase in net plant rate (**Figure 3F**).

**Figure 3.**
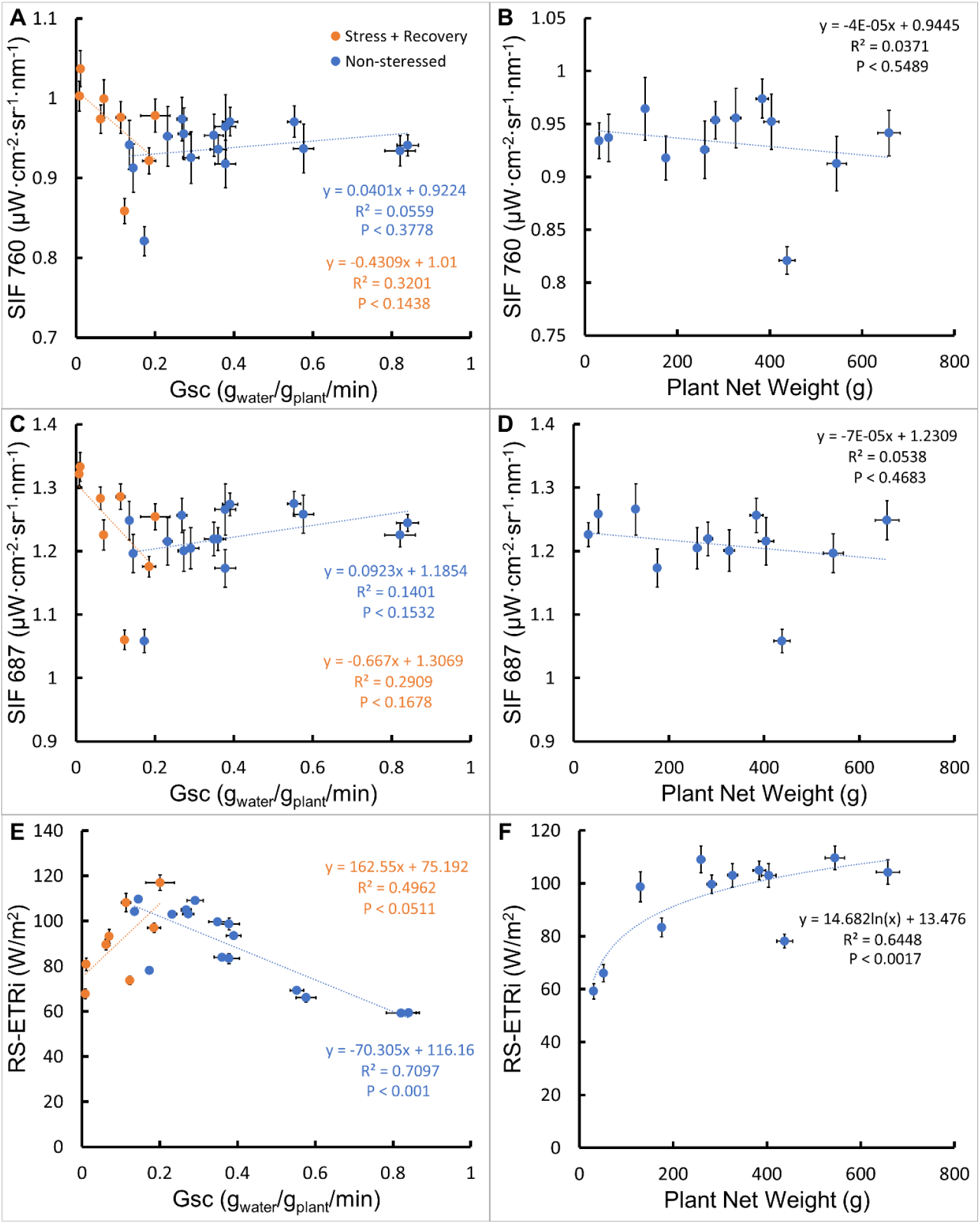
Correlations between SIF based indices and high-throughput platform parameters. Panels **(A), (C), (E)**represent correlations of SIF_760_, SIF_6S_7 and RS-ETRi with canopy conductance (Gsc) and Panels **(B), (D), (F)**with net plant weight. DAT = Days after transplanting. 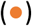 Stress + Recovery – values from DAT 18 – DAT 31, values are means of 46-52 biological replicates ± SE. 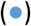 Non-stressed - values from DAT 5 – DAT 14, values are means of 20–52 biological replicates ± SE. Panels A–C, are means of 66-72 biological replicates ± SE. Panels D-F are means of 18–20 biological replicates ± SE.

### SIF based indices as predictors for drought stress

We were interested to determine if SIF based indices can perform as early indicators of drought stress, compared with traditional reflectance VIs and platform gravimetric parameters. Tomato plants were exposed to a stress period with recovery **(Figure 4)**. Values of the drought-treated plants were normalized with the average value of the control-group plants measured at the same time, in order to create a similar scale between all the plants.

**Figure 4.**
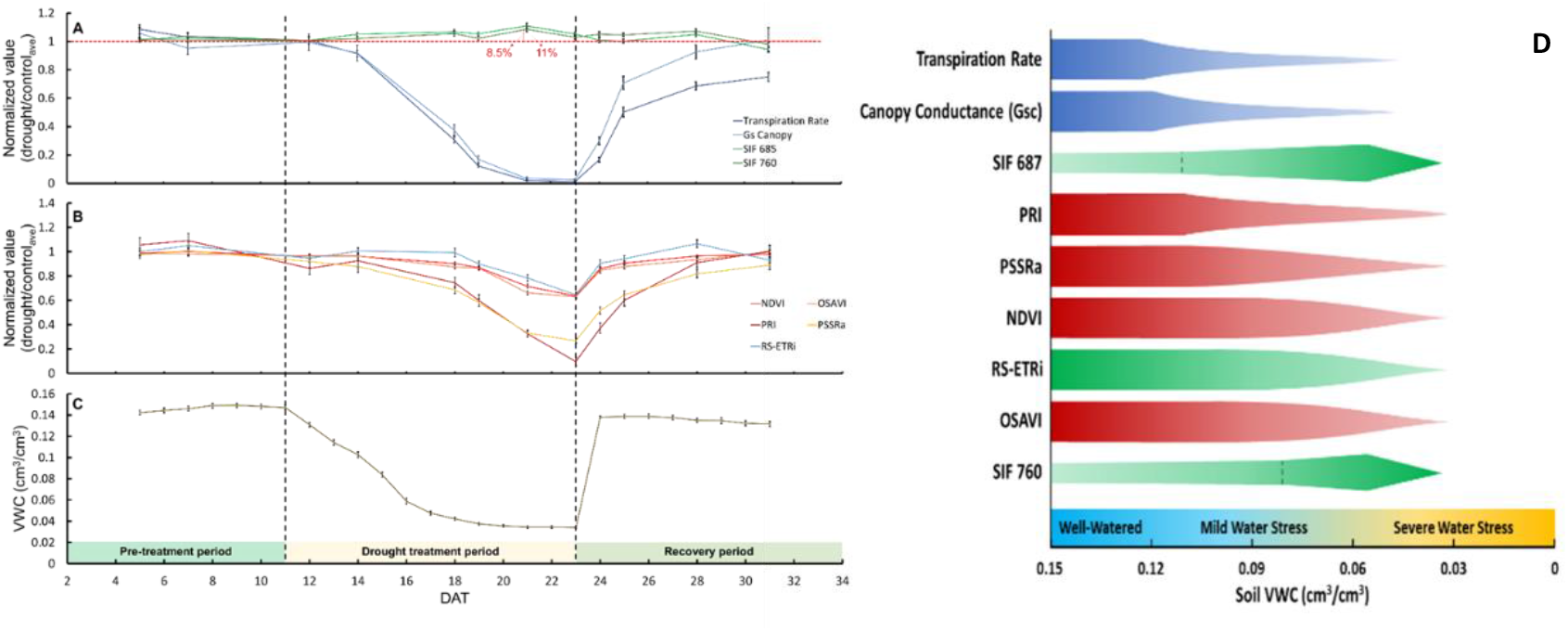
Temporal response of transpiration, SIF based indices, and VIs to drought and recovery periods. (**A**) Normalized values of SIF_687_ 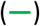, SIF_760_ 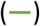, average transpiration rate at noon 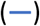 and average canopy conductance 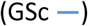. The red dashed line indicates that there is no difference between the drought and control groups (value of 1). (**B**) Normalized values of NDVI 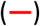, OSAVI 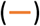, PRI 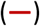, PSSRa 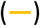 and RS-ETRi 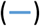. Values are means of 52 biological replicates ± SE. (**C**) Volumetric water content of the soil at noon throughout the experimental period. (**D**) Diagram of early response of each parameter analyzed. Thickness determines value. Thinner bulb means decreased value of a parameter. Values are means of 40 replicates ± SE.

High-throughput platforms parameters canopy conductance and transpiration rate track accurately the actual soil moisture content recorded by the system **(Figure 4A, 4C)**. SIF_687_ and SIF_760_ increased by up to 11% and 8.5%, respectively, among the drought-treated plants, as compared to the well-watered control plants **(Figure 4A, green curves)**. On DAT 23, the last and most severe day of the drought, SIF values decreased a bit and continued to decrease during the recovery period **(Figure 4A Green lines)**. PRI and PSSRa were first to respond to the stress (**Figure 4B, start of stress period)**, though not significantly, both in time and effect. NDVI and PRI responded only 3 days after the stress period started (DAT 14), and RS-ETRi responded only 7 days later towards the maximum stress of the plant (DAT 18).

### SIF based indices as screen predictors between introgression lines

SIF based indices as well as traditional VIs were checked for their ability to identify differences between different genotypic cultivars of tomato (*S. Lycopersicum*). Four Introgression Lines (IL) of tomato were selected: IL-2-6-5, IL 5-2, IL 6-4, IL 8-1-1 (**Table 1)**. Both gravimetric parameters and remote sensing indices did not show statistically significant differences between the four introgression lines.

**Table 1.** Characterization of well-watered tomato plants from different introgression lines 12 days after transplanting (DAT), as expressed by gravimetric measurements-daily transpiration and canopy conductance; and remote-sensing indices-traditional vegetation indices and SIF based indices. *n* = 10–15 plants in each group.

| Parameter | Introgression Lines |  |  |  |
| --- | --- | --- | --- | --- |
|  | IL 2-6-5 | IL 5-2 | IL 6-4 | IL 8-1-1 |
| Daily Transpiration | 484.0 ± 26.21 | 504.6 ± 26.26 | 535.6 ± 18.55 <sup>†</sup> | 428.4 ± 23.57 <sup>†</sup> |
| Gsc | 0.37 ± 0.02 | 0.39 ± 0.01 | 0.38 ± 0.01 | 0.34 ± 0.01 |
| NDVI | 0.68 ± 0.02 <sup>†</sup> | 0.80 ± 0.01 <sup>†</sup> | 0.73 ± 0.02 | 0.76 ± 0.01 |
| MTVI2 | 0.25 ± 0.01 | 0.29 ± 0.01 | 0.27 ± 0.01 | 0.29 ± 0.00 |
| PRI | -0.02 ± 0.003 <sup>†</sup> | -0.04 ± 0.002 <sup>†</sup> | -0.03 ± 0.004 | -0.03 ± 0.002 |
| PSSRa | 5.96 ± 0.58 | 8.34 ± 0.78 | 7.75 ± 0.86 | 7.78 ± 0.50 |
| SIF<sub>687</sub> | 1.22 ± 0.03 | 1.33 ± 0.03 | 1.28 ± 0.03 | 1.24 ± 0.04 |
| SIF<sub>760</sub> | 0.93 ± 0.02 | 1.02 ± 0.02 | 0.98 ± 0.02 | 0.95 ± 0.03 |
| RS-ETRI | 82.65 ± 4.98 | 104.4 ± 5.41 | 97.12 ± 5.10 | 96.10 ± 5.54 |

<sup>†</sup> Significant difference between groups at $p < 0.05$

Experimenting with drought, IL 5-2 and IL 8-1-1 show differences in fresh, dry biomass and cumulative transpiration (CT) for both the control and the drought treatments, but no differences in Water Use Efficiency **(Table 2)**. (We selected these two representatives due to the biggest differences in response to drought between the two)

**Table 2.** Comparison of physiological characteristics of the tomato IL lines IL5-2 and IL8-1-1 under well-watered (control) conditions and drought conditions. All measurements were performed at the end of the experimental period. n_control_ = 4, n_drought_ = 11.

| Lines | Dry weight (g) |  | CT (g/1000) |  | Shoot fresh weight (g) |  | Water Use Efficiency<br>(g/ml)<br>* 1000 |  |
| --- | --- | --- | --- | --- | --- | --- | --- | --- |
|  | Control | Stress | Control | Stress | Control | Stress | Control | Stress |
| IL 5-2 | 87.3 ± 4 | 40.9 ± 2 | 18.7 ± 0.3 | 9.6 ± 0.4 | 658 ± 45 | 315 ± 15 | 4.4 ± 0.13 | 4.0 ± 0.04 |
| IL 8-1-1 | 72.1 ± 5 | 31.3 ± 1 | 16.1 ± 0.8 | 7.1 ± 0.4 | 546 ± 21 | 233 ± 14 | 4.2 ± 0.10 | 4.1 ± 0.08 |

\* $P < 0.05$

We also examined the responses of the lines to drought stress and found that although IL5-2 suffered from a higher penalty during the drought stress, it recovered more rapidly and more efficiently than IL8-1-1 (**Supplementary Table S4**). The two lines showed different behavior in transpiration during well-watered conditions but they did not show any significant differences when checked with VIs and SIF based indices **(Supplementary Figure 4).**However, when the two lines were exposed to drought conditions, the drought-treated IL5-2 plants exhibited higher DT and Gsc during the entire recovery period and on some days during the pre-treatment period; whereas IL8-1-1 had a higher Gsc on DAT 19, during the drought period (**Figure 5A, 5C**). IL5-2 had higher NDVI values on the first measurement day, DAT 5, and at the end of the recovery period (DAT 28 and 31; **Figure 5H**). IL5-2 also had significantly lower PRI on DAT 5, but not after that date (**Figure 5G**). No significant differences were observed between the groups in terms of SIF, RS-ETRi or calculated plant weight (**Figure 5B, 5D–5F**).

**Figure 5.**
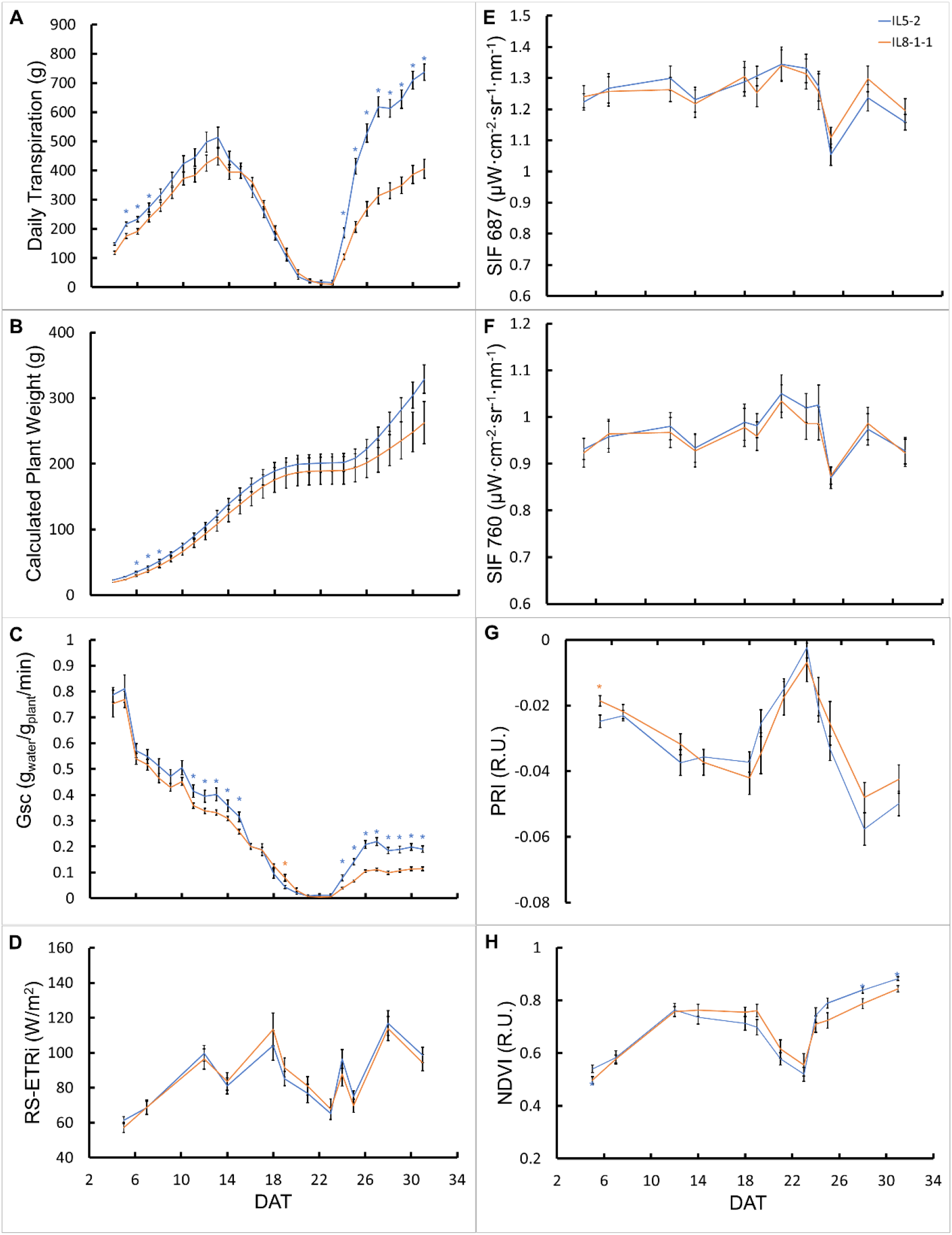
Analysis of VIs, SIF based indices and high through-put parameters throughout the experimental period in tomato (*S. Licopersicum*) IL5-2 and IL8-1-1 exposed to the drought treatment. IL5-2 and IL8-1-1 were compared in terms of (**A–C**) physiological traits measured gravimetrically by the FPP, (**D–F**) SIF and RS-ETRi and (**G–H**) selected VIs. Each point is a mean of 10–11 biological replicates. An asterisk (*) indicates a significant difference between groups (*t*-test, *P* < 0.05). In each panel, the color of the asterisk corresponds to the group that had the significantly higher value.

## Discussion

The performance of high-frequency measurements, high-throughput photosynthesis analysis is a challenge for which conventional tools are not yet able to provide a satisfactory solution (Du et al., 2020; Fu et al., 2022). In the current study, Vegetation and SIF based indices were used in order to overcome this challenge. DCMU blocks the electron transport chain between quinone a and quinone b site and by that induces an abrupt increase in fluorescence emission (Ridley, 1977). (Pinto et al., 2016) proved that spraying DCMU on wheat and maize leaves increase their SIF signal. These studies corroborate our findings **(Figure 1)** and by that prove that the construction of the spectroradiometer on top of the high-throughput phenomics system including the calculation that extracts and quantifies the signal are correct. SIF values decreased together with PRI values on the background of an increase in chlorophyll a and b content **(Figure 2)**. The decrease in the SIF_687_ is expected, because as chlorophyll concentration increased, red-SIF is re-absorbed back into the canopy (Van Der Tol et al., 2009). A decrease in the SIF_760_ with increase in chlorophyll content is also possible due the fact that increase in chlorophyll content implies for an increase in canopy size, which prevents the escape of SIF_760_ back to the environment (Guanter et al., 2014). We interpret the decrease in the PRI values with increase in the chlorophyll content as a measure of health of the plant, where more violaxanthin is generated in order to harvest more light when more photosystems are synthesized (Gamon et al., 1992).

There are increasing evidences in literature for the complex relationship between canopy conductance and SIF based indices. (Shan et al., 2019) shows promising new evidence for SIF signals extracted from satellite to corelate positively with canopy conductance of wheat and to some extent trees and grasses. In our hands, SIF_760_ and SIF_687_ showed negative relationship with canopy conductance during stress **(Figure 3A, 3C)**, and Electron transport rate index showed positive correlation canopy conductance on the background of stressed plants **(Figure 3E)**. During drought, stomata are closed and reduction in photosynthetic activity (Reddy et al., 2004), and it is therefore only natural that absorbed energy will be dissipated either by NPQ or emission of fluorescence (Baker, 2008), both corroborating our result.

Additional to this, although many studies have reported on a strong relationship between SIF and GPP (Guan et al., 2016; Guanter et al., 2014), we did not find meaningful correlations between plant biomass gain and SIF_687_ or SIF_760_ (**Figure 3B, 3D**). There was significant positive (logarithmic) correlation between RS-ETRi and the plant weight measured by the FPP (**Figure 3F**). This is expected in view of the logarithmic correlation between the special SIF term within this index (Liran, 2022) which implies for complex relationship between light use efficiency and biomass production.

SIF is directly related to photosynthetic activity in plants as it is the surplus energy which is emitted back to the environment and not used (Baker, 2008). Therefore, it is regarded as a more direct probe of carbon fixation than reflectivity-based measurements (Meroni et al., 2008). We therefore hypothesized that SIF would be a more useful remote-sensing indicator for drought stress than traditional VIs (Xu et al., 2021). Normalized SIF_687_ and SIF_760_ increased in response to the water stress **(Figure 4)**. (Jonard et al., 2020) suggested that there are three phases of the relationship between SIF and drought stress. In the first phase (before stress), a rise in incoming radiation leads to an increase in SIF and in photosynthesis, until all of the chlorophyll/rubisco molecules are saturated (Wieneke et al., 2018). In the second phase (mild stress), when soil water availability becomes a limiting factor for plant performance, the photosynthetic rate decreases due to stomatal and non-stomatal limitations. This decrease in photosynthesis is linked to an increase in NPQ (Frankenberg & Berry, 2018). Our measurements, which were made on a single plant canopy, revealed increases in SIF_687_ and SIF_760_ in response to water stress, followed by a decrease on the last day of the stress (**Figure 4A**), which was also the most severe. These results are in line with the three-phases theory and are similar to the findings of Wohlfahrt et al. (2018) regarding the response of SIF to water stress. The whole-plant transpiration rate and stomatal conductance showed stronger and faster reactions to drought than SIF (**Figure 4A**). Similar results were reported by Helm et al. (2020), who measured stomatal conductance and photosynthesis in terms of gas exchange simultaneously with SIF over a similar period of time. The SIF-based index, RS-ETRi, responded to drought in a similar way to NDVI and OSAVI (**Figure 4B**).

Ihuoma & Madramootoo (2017) relate reflectance VIs to plant health and plant water stress. However, the temporal responses of the four VIs were not significantly different from one another, although the PSSRa and PRI indices responded more quickly than the others (**Figure 4B**). The earlier decrease in PSSRa might be a result of an increase the catalytic activity of chlorophylls and degradation of photosynthetic pigments (Enneb et al., 2020). The decrease in Chl content is a commonly observed phenomenon under drought stress (Ashraf & Harris, 2013). Besides its earlier response, PRI also showed a much stronger reaction than the other three indices at the same time and soil water content (**Figures 4B**). This observation is reasonable as PRI is considered to be a good indicator of the level of plant water stress (Thenot et al., 2010). It is very sensitive to heat dissipation, which increases under water-stress conditions (Panigada et al., 2014), and to the rapid changes in carotenoids that occur through the de-epoxidation of the xanthophyll pigments (Gamon et al., 1992). In contrast, the traditional greenness indices NDVI and OSAVI sense the stress only when the plant structure begins to be affected (Panigada et al., 2014).

Finally, it has been suggested that remote sensing VIs can report on biodiversity of plants (Wang & Gamon, 2019). We tested whether SIF based indices together with traditional VIs can distinguish between four Introgression Lines (ILs) of tomato (*S. Licopersicum*). The selected lines were chosen according to their physiological characteristics, as well as a yield characterization performed by Gosa et al. (2022). We decided to examine the lines that were most different from one another. Line IL5-2 was classified as an ‘ideotype’, and line IL8-1-1 was classified as a ‘survival’ phenotype by Gosa et al. (2022) and their yields under field and greenhouse conditions were reported to be significantly different. The higher biomass and transpiration levels of IL5-2 (Supplementary Table S3) are not surprising, since previous studies also reported higher biomass and fruit yield among IL5-2, as compared with IL8-1-1, under both well-watered and drought conditions (Gosa et al., 2022). IL5-2’s anisohydric-like nature led to increased transpiration during recovery from stress; whereas its biomass appeared to increase relatively slowly, which decreased its Water Use Efficiency (WUE). The survival line, IL8-1-1, exhibited a more isohydric strategy, decreasing its transpiration at a slightly slower rate during stress, but not fully regaining it during the recovery period. Neither of the tested remote-sensing methods, including SIF, has proved to be as sensitive or accurate as the gravimetric system in distinguishing the tomato ILs. It is possible that the SIF and VI measurements cannot discern subtle physiological differences between plants when measuring a single canopy at a time. Furthermore, research into diurnal and inter-day changes in SIF has shown that SIF measured across days is affected mainly by chlorophyll content; whereas diurnal changes in SIF reflect photosynthetic activity (Mohammed et al., 2019). Our results indicate that SIF measured once a day, on a single plant canopy, does not provide an advantage over gravimetric measurements or other commonly used methods for detecting physiological differences between plants.

There are two main limitations to this study. First, we did not validate photosynthesis measurements on leaf level with established hand held fluorometer or gas exchange systems. Liran et al. (2020) managed to show that electron transport rate measured at leaf level is correlated with RS-ETRi index in greenhouse conditions and C3 plants, but this examination was performed in a unique set-up which prevented the plant to reach high level of NPQ and photoinhibition. On the other hand the nature of the high-throughput system to measure high number of samples in a short period of time a tedious task, and therefore the authors suggest to at least add the dimension of effective quantum yield measurements which are fast enough to perform in future experiment. Second, measurements were performed only during a distinct time range – early noon, as suggested by the study of Prior et al. (1997). Measuring stress along different parts of the day may shed a light on the carbon balance and how plants cope with stress along a diel cycle. Finally, One of the biggest challenge humanity face is the need in new crop breeds that can sustain extreme events of droughts and weather conditions (Tester M, 2010). This conjugated system will be able to assist and screen for a better crop cultivar with optimal physiological as well as photosynthetic characteristics.

## Material and Methods

### Area of study

The study was conducted between May 2019 and June 2021 at the Robert H. Smith Faculty of Agriculture, Food and Environment of the Hebrew University of Jerusalem, in Rehovot, Israel. The experiments for this study were carried out in the iCORE functional-phenotyping greenhouse (https://plantscience.agri.huji.ac.il/icore-center). The Photosynthetically active radiation (PAR), temperature (T), vapor pressure deficit (VPD) and relative humidity (RH) measured during the experimental period were continuously monitored (**Supplementary Figure 1**).

### Devices and systems

The setup and measurement techniques of the system are described thoroughly in Halperin et al. (2017) and Dalal et al. (2020). Essentially, a thoroughly multi sensor system is able to record soil moisture, water incoming and outgoing from each pot, and it is situated over a scale.

### Plant material and treatments

Tomato (*S. lycopersicum*) introgression lines (Ils) developed by crossing the M82 cultivar and the wild-type *S. pennellii* were used (supplementary S4). Each of the lines contained a single homozygous restriction fragment-length polymorphism of a *S. pennellii* chromosome segment (Eshed & Zamir, 1995). The seeds were kindly provided by Prof. Dani Zamir of the Hebrew University of Jerusalem. Based on previous work, we selected four representative lines from the Il population for use in the main experiment. The lines differed in their biomass and yield under well-irrigated conditions, as well as their drought tolerance and resilience (Gosa et al., 2022).

### Acquisition of spectral data

We took samples with the spectrometers from all 72 plants in the experimental array at midday (between 12:00 to 13:00), 12 times over the course of the experimental period. The time of day was selected to match the time of maximum transpiration of the plants. Overall, there were two measuring points during the pre-treatment phase, six measuring points during the drought phase and four measuring points during the recovery phase. To measure all 72 plants in a short time, we used a polypropylene cart with a custom-made adjustable arm on which we placed the two spectrometers. The arm allowed us to raise and lower the spectrometers as needed. To measure SIF and other VIs by remote sensing, we used two sets of custom high-sensitivity VIS-NIR spectrometers (Maya 2000 Pro, Ocean Insight, Orlando, FL, USA) calibrated with a NIST-standardized halogen light (HL-2000-LL, Ocean Insight, Orlando, FL, USA) with a slit size of 5 μm and grating of 600 lines/mm. The wavelength range of the spectrometer was 400–838 nm and the spectral resolution was 0.32 nm (2064 values in each acquisition). The spectrometer was coupled with an optical fiber with a core diameter of 400 μm. The high sensitivity of the spectrometer was necessary to retrieve the SIF and to calculate reflectance-based VIs. The spectrometers were recalibrated once a year with a suitable light source for VIS-NIR (HL-2000-HP, Ocean Insight). One spectrometer was aimed upward to measure the downwelling irradiance from the sun (diffuse plus direct beam radiation). To do that, the optical fiber was coupled with a cosine corrector optical diffuser (CC-S-DIFFUSE Spectralon Diffuser, Labsphere, Inc., NH, USA) to expand the field of view to 180º. The second spectrometer was aimed downward to measure the upwelling radiance reflected from the canopy of the plant. This fiber was bare. This system allowed us to scan all of the plants in about 30 min. The optical fiber was placed at a distance of about 20–40 cm from the canopy, which allowed the capture of a circular area about 8.8–17.6 cm in diameter, which suited the center of a single plant canopy. To verify that the background of the captured canopy was not affecting the spectrum, we captured several spectra of the background only, without any plants (plastic pot, EVA surface cover, tubes). The background spectra were compared with the plant spectra and no significant effect was found (**Supplementary Figure S2**).

### Measurement of chlorophyll concentrations

Chlorophyll concentrations in leaf samples were measured three times during the experimental period, once during each of the three different phases of the experiment: pre-treatment, drought and recovery. A chlorophyll-extraction protocol with DMSO (>99.7%, Thermo Fisher Scientific, Waltham, MA) was used to calculate chlorophyll concentrations. The extraction protocol included collecting five 0.5-cm discs from the youngest mature leaf of each plant and storing the collected tissue at −76°C for at least 24 h (Moran, 1982). Each frozen sample was then combined with 2 mL DMSO and incubated in 50°C water for 2 h (Novák et al., 2013). After incubation, the absorbance spectrum between 400 to 800 nm was measured by a multimode microplate reader (Tecan Spark, Tecan, Switzerland) for 200 μL of each sample. Concentrations of chlorophyll a and chlorophyll b were calculated based on the equations described by Wellburn (1994).

### Course of the experiment

The experiment was designed to include three periods:

1. A **pre-treatment** took 9 days period during which all of plants were well-irrigated, receiving seven pulses of water during the night. During that phase, the FPP calculated the water-use efficiency of the plants, a value that was then used it to estimate the weight of the plants in real time during the drought treatment. six pulses during the nighttime (between 20:30 and 02:30) only, to allow the proper measurement of daily transpiration
2. A **drought** treatment period during which the drought-treated plants were exposed to a gradual reduction of soil water content with a fixed dehydration rate of 70–90 g per day, until transpiration was stopped entirely for 2 days and the plants wilted.
3. A **recovery** period of 11 days, during which the drought-treated plants recovered and were well-irrigated through the end of the experiment. the drought-treated plants received increased irrigation with 10 pulses of 2–4 min each. Starting the next day, they received the same irrigation as the control-group plants.

### Extraction of spectral data and calculation methods

Raw data from the spectrometer is obtained in terms of photon counts per wavelength and needs to be converted into light flux units (μW*cm^-2^*sr^-1^*nm^-1^). We converted it according to Liran (2020) (supplementary S9). SIF was calculated using the iFLD method, as dictated by Alonso et al.(2008).

### Calculation of vegetation indices (VIs)

VIs are mathematical combinations of two or more spectral wavelengths. Various indices can be calculated based on the spectral data acquired with our spectrometers (400–838 nm). In this study, only some of the indices we calculated were examined and analyzed (**Table 1**).

**Table 1.**
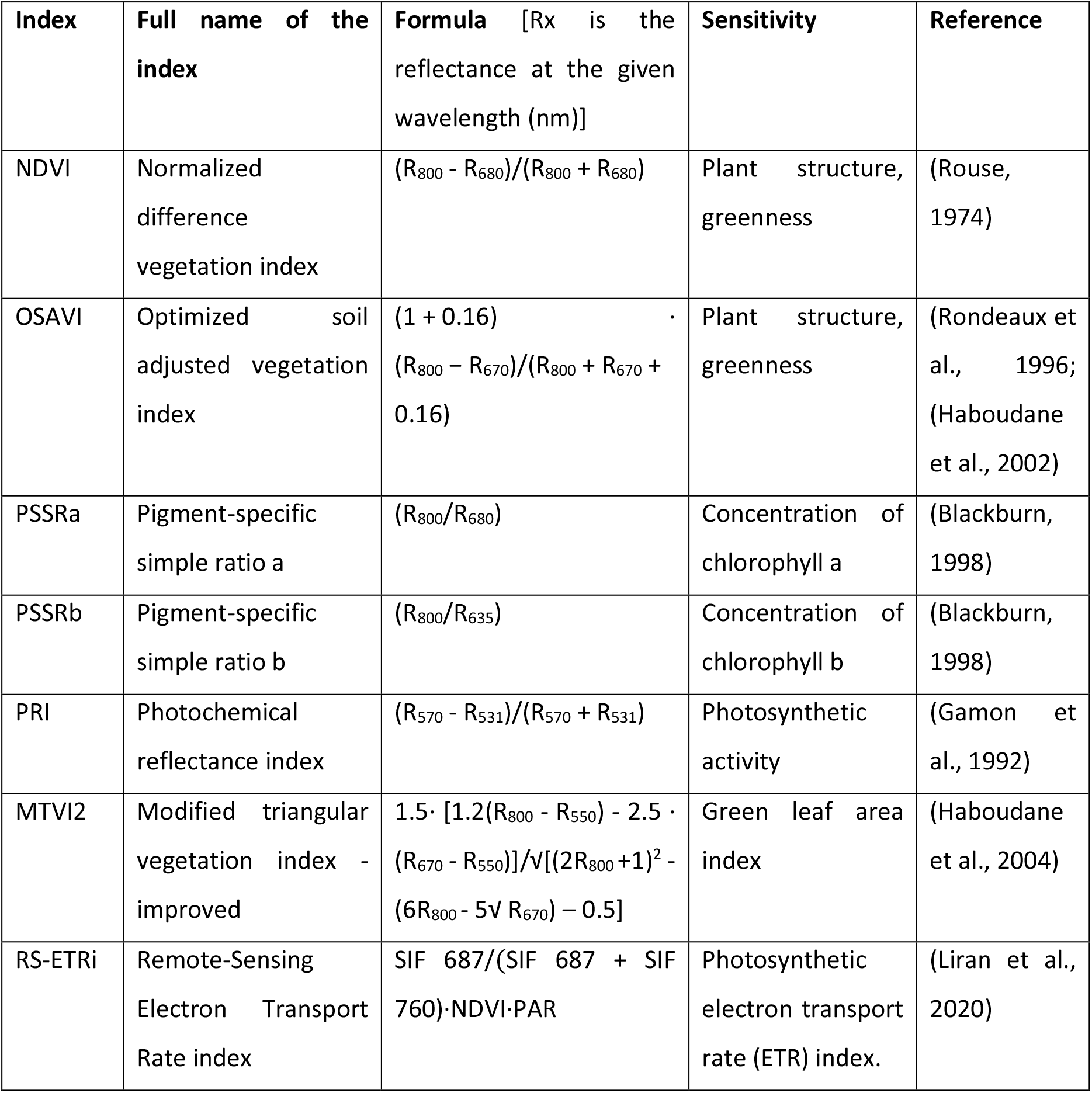
Spectral vegetation indices used in this study.

### Statistical analysis

The data analysis was performed using the JMP ver. 15.0 (SAS Institute INC., NC, USA) statistical package and Excel (Microsoft, WA, USA). In all of the comparison tests, groups were checked for normal distribution with the Shapiro–Wilk’s test and the homogeneity of variance was examined with Levene’s test. If both tests were satisfied, *t*-tests were used to compare two groups and analysis of variance (ANOVA) followed by Tukey’s honestly significant difference (HSD) was used for multiple comparisons. When comparing to a control group, Dunnett’s test was used. If the normality criteria were violated, Wilcoxon/Kruskal–Wallis’s nonparametric ANOVA was used. All tests were performed at a significance level of α< 0.05. The mean values for all results are presented with ± S.E. The squared values of Pearson correlation coefficients were also calculated in regression analyses. The graphs were plotted using Microsoft Excel 2019.

## Acknowledgments

This research was supported by ISF (the Israel Science Foundation), Grant No. 1043/20 and ISF-NSFC Grant No. 2436/18.

## Notes

### Competing Interest Statement

The authors have declared no competing interest.

